# Robustness of the Dorsal morphogen gradient with respect to morphogen dosage

**DOI:** 10.1101/739292

**Authors:** Hadel Al Asafen, Prasad U. Bandodkar, Sophia Carrell-Noel, Gregory T. Reeves

## Abstract

In multicellular organisms, the timing and placement of gene expression in a developing tissue assigns the fate of each cell in the embryo in order for a uniform field of cells to differentiate into a reproducible pattern of organs and tissues. This positional information is often achieved through the action of spatial gradients of morphogens. Spatial patterns of gene expression are paradoxically robust to variations in morphogen dosage, given that, by definition, gene expression must be sensitive to morphogen concentration. In this work we investigate the robustness of the Dorsal/NF-κB signaling module with respect to perturbations to the dosage of maternally-expressed *dorsal* mRNA. The Dorsal morphogen gradient patterns the dorsal-ventral axis of the early *Drosophila* embryo, and we found that an empirical description of the Dorsal gradient is highly sensitive to maternal *dorsal* dosage. In contrast, we found experimentally that gene expression patterns are highly robust. Although the components of this signaling module have been characterized in detail, how their function is integrated to produce robust gene expression patterns to variations in the *dorsal* maternal dosage is still unclear. Therefore, we analyzed a mechanistic model of the Dorsal signaling module and found that Cactus, a cytoplasmic inhibitor for Dorsal, must be present in the nucleus for the system to be robust. Furthermore, active Toll, the receptor that dissociates Cactus from Dorsal, must be saturated. Finally, the vast majority of robust descriptions of the system require facilitated diffusion of Dorsal by Cactus. Each of these three recently-discovered mechanisms of the Dorsal module are critical for robustness. Our work highlights the need for quantitative understanding of biophysical mechanisms of morphogen gradients in order to understand emergent phenotypes, such as robustness.

**Author Summary:** The early stages of development of an embryo are crucial for laying the foundation of the body plan. The blueprint of this plan is encoded in long-range spatial protein gradients called morphogens. This positional information is then interpreted by nuclei that begin to differentiate by expressing different genes. In fruit fly embryos, the Dorsal morphogen forms a gradient along the dorsal-ventral axis, with a maximum at the ventral midline. This gradient, and the resulting gene expression patterns are extraordinarily robust to variations in developmental conditions, even during early stages of development. Since positional information is interpreted in terms of concentration of the morphogen, one would expect that doubling or halving dosage would result in disastrous consequences for the embryo. However, we observed that development remains robust. We quantified the effect of dosage by experimentally measuring the boundaries of 2 genes, - *sna* and *sog*, expressed along the DV axis and found that variation in the boundaries of these genes was minimal, across embryos with different dosages of Dl. We then used a mathematical model to discern components of the Dl system responsible for buffering the effects of dosage and found three specific mechanisms – deconvolution, Toll saturation and shuttling

## Introduction

The morphogen concept forms the basis of many models of developing tissues. Through their concentration gradients in space, morphogens send positional information to cells and direct them to develop in specific ways depending on their location within a tissue. The roles of these signals range from the development of the initial polarities of embryos to specification of cell identity in specific tissues, and the nervous system in both vertebrates and *Drosophila* [1]. Tissue patterning is often initiated by the cells’ concentration-dependent response to the morphogen gradient: cells throughout the tissue are subject to different levels of morphogen, depending on their position within the field, and accordingly, express distinct target genes. Thus, the quantitative shape of the morphogen gradient is critical for patterning, with cell-fate boundaries established at specific concentration thresholds. The cells’ sensitivity to morphogen concentration also implies that any shift in the morphogen distribution is expected to result in an accompanying shift in patterning. Therefore, perturbations to the morphogen dosage or production rate, which should change the morphogen distribution, should in turn perturb gene expression patterns.

Indeed, early models of morphogen gradient formation assumed the gradient scaled globally with the morphogen dose (*e*.*g*., when one copy of the gene encoding the morphogen is lost, the entire distribution is divided by two). Such “dosage-scaling” models predicted that catastrophic shifts in target gene expression domains would occur when the dose of morphogen is altered [2, 3]. In contrast, experimental observations have shown that the spatial positioning of morphogen target genes shift only minimally when morphogen dosage is perturbed [2–5], with some notable exceptions [6, 7]. Thus, there exists a paradox between the sensitivity of cells to morphogen concentration and the robustness of tissue patterns with respect to morphogen dose, which implies a mechanism that prevents robust morphogen gradient systems from scaling with morphogen dose. One such mechanism is self-enhanced ligand degradation, where the ligand (morphogen) upregulates its own inhibitor, and which has been suggested to explain experimentally-observed robustness [3,4,8,9]. However, this mechanism does not apply to all morphogen gradient systems. In particular, the Dorsal/NF-κB signaling network in *Drosophila* embryos does not clearly exhibit the self-enhanced degradation mechanism.

The NF-κB module, conserved from flies to humans, is implicated in several cellular responses/phenotypes, including tissue patterning, inflammation, innate immunity, proliferation/apoptosis, and cancer [10–14]. The maternal transcription factor Dorsal (Dl), homologous to mammalian NF-κB, patterns the dorsal-ventral (DV) axis of the developing *Drosophila melanogaster* embryo to specify mesoderm, neurogenic ectoderm, and dorsal ectoderm cell fates (Roth et al., 1989; reviewed in [16–18]. In the early embryo, Dl protein is initially uniformly distributed around the DV axis.

During nuclear cleavage cycle (nc) 10, the nuclei migrate to the periphery of the syncytial blastoderm and the Dl gradient begins to be established. The IκB homolog Cactus (Cact), which is also maternally-supplied, binds to Dorsal, retaining it outside the nuclei. Toll, the *Drosophila* homolog of the Interleukin 1 receptor, is active on the ventral side of the embryo, where it signals through Pelle kinase to phosphorylate the Dl/Cact complex [19], which results in dissociation of Dl from Cact, allowing Dl to enter the nuclei, where it regulates gene expression. Because Toll signaling is spatially asymmetric, a nuclear gradient of Dl forms, with a peak at the ventral midline and a Gaussian-like decay in space to become nearly flat at approximately 45% of the embryo’s circumference (**Figure 1A**) [5, 20]. From 45% to 100% ventral-to-dorsal coordinate, the gradient has a shallow downward slope to achieve non-zero basal levels at the dorsal midline [5,20,21]. Our computational studies have suggested the non-zero basal levels are primarily composed of Dl/Cact complex in the dorsal-most nuclei, not free Dl [22].

**Figure 1:**
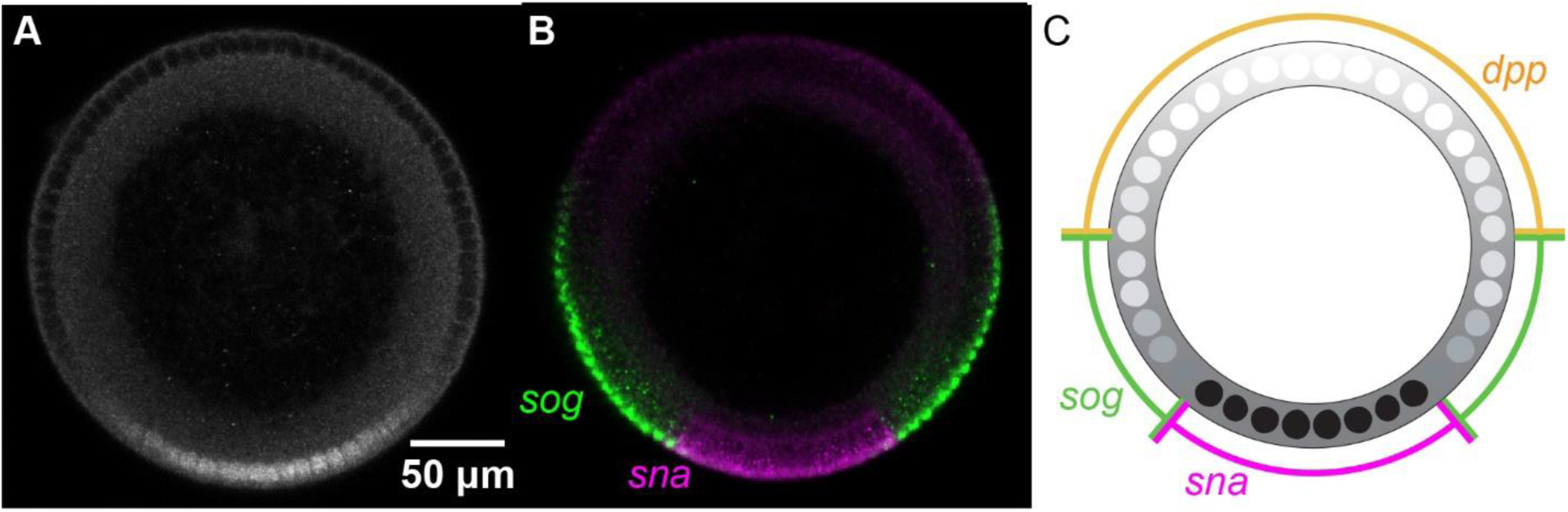
The protein Dorsal patterns the DV axis of the *Drosophila* embryo. (A) An antibody staining against Dorsal in an NC 14 embryo. (B) mRNA expression of Dorsal target genes *sna* (magenta) and *sog* (green). (C) Illustration of the borders of gene expression. We use these borders to quantify and compare the extent of domain of dl target genes. Embryo cross-sections are oriented so that ventral is down.

As shown in **Figure 1B**, different genes are turned on at different concentrations of Dl [16, 23]. It can be both an activator and a repressor of transcription. At high concentrations of Dl on the ventral side of the embryo, high threshold genes such as *snail* (*sna*) are expressed. In the lateral part of the embryo, intermediate Dl levels activate the expression of low threshold genes such as short gastrulation (*sog*). The domains of these genes can be quantified using measurements of the dorsal border and ventral border (**Fig 1C**).

While the copy number of maternal *dl* has been shown to affect the Dl gradient and downstream tissue structure, the phenotypes are subtle. Embryos from mothers heterozygous for a *dl* null allele (1x *dl*) have shorter, wider, and flatter Dl gradients as compared to wildtype [5,24–26]. While these embryos have a weakly dorsalized phenotype, female flies with a half dose of *dl* produce a high fraction of viable progeny at room temperature [27, 28]. Furthermore, measurements in a handful of embryos (n < 12) found no statistical shift in the *sog* expression domain [5], and a shift of roughly only one cell diameter in the *sna* domain [26]. The altered shape of the Dl gradient has recently been attributed to a combination of two novel observations. First, Cact acts to facilitate the diffusion of Dl (i.e., “shuttling” of Dl by Cact), which results in a net flux of Dl to the ventral side of the embryo [26]. And second, active Toll receptor complexes are saturated by Dl/Cact complex [26]. Together, these processes act to accumulate Dl on the ventral side in wildtype embryos, but accumulate Dl in ventral-lateral regions in 1x *dl* embryos. Furthermore, experimental evidence strongly suggests the shuttling mechanism is required for viability of 1x *dl* embryos, as embryos from heterozygous *dl* mothers that also have compromised shuttling are non-viable [26].

In a similar manner, embryos with overexpression of excess, transgenic copies of *dl* (4x *dl*) are only weakly ventralized, and a large fraction still hatch [29]. Given the subtlety of the 1x and 4x *dl* phenotypes, and the viability of the embryos, one may ask whether this implies the Dl gradient system is robust, and if so, whether the robustness requires special mechanisms, such as shuttling and Toll saturation [26]. As mentioned above, dosage-scaling models are typically sensitive to dosage; however, such a model of the Dl gradient has not been analyzed for robustness of gene expression with respect to variations in morphogen dosage.

In this work, we used empirical and computational modeling, together with quantitative measurements of the Dl gradient and domains of target gene expression, to investigate the robustness of the Dl gradient system with respect to dosage of maternal *dl*. First, we showed that a dosage-scaling formulation of the Dl gradient has an unacceptably high sensitivity to the maternal dosage of *dl*, even in the best-case scenario, in which basal levels are composed primarily of Dl/Cact complex and there is negligible Dl activity at the dorsal midline [22]. In particular, in the absence of a mechanism to prevent dosage-scaling, doubling or halving the maternal *dl* dosage is predicted to result in drastic perturbations to gene expression. Next, we experimentally measured gene expression domains and the Dl gradient width in embryos from mothers with *dl* dosages of 1x (heterozygous null for maternal *dl*), 2x (wildtype), and 4x (expressing two copies of a *dl* rescue construct; Carrell et al., 2017; Reeves et al., 2012) and showed that, in contrast to the predictions of the dosage-scaling model, the perturbations to patterns are minimal. To identify the possible mechanism for this robustness, we analyzed a computational model of the Dl/Cact system. Our model is based on previously published models in which Dl and Cact can interact, enter the nuclei, and diffuse between “cytoplasmic compartments” surrounding the nuclei [21,22,24,26,30]. The active Toll signaling complex, which is limited to the ventral side of the embryo, acts as a Michaelis-Menten-like enzyme to favor dissociation of the Dl/Cact complex. We constrained the model using our measurements of Dl target gene expression in 1x, 2x, and 4x embryos. A random parameter search showed that several parameter sets resulted in robust phenotypes. Our analysis of the robust parameter sets showed that robustness can rarely be achieved unless (1) the free Dl nuclear levels drop to near zero on the dorsal side of the embryo [22], (2) significant facilitated diffusion by Cact occurs [26], and (3) active Toll signaling can be saturated by Dl/Cact complex [26]. Quantitative analysis can be used to assess rigorously the robustness of different patterning models. Applying the same modelling principles to other systems might identify additional mechanisms that underlie robust patterning by morphogen gradients in development.

## Results

### Sensitivity of a dosage-scaling model of the Dl gradient

Early models of morphogen gradients exhibited “dosage-scaling,” in that these descriptions of the morphogen gradient scaled globally, in a multiplicative manner, with morphogen dosage. Morphogen gradients predicted by these models were highly sensitive with respect to morphogen dosage [3,31,32]. However, these models focused on exponential-like morphogen distributions, whereas the Dl gradient is Gaussian-shaped [5,20,26]. Therefore, to determine the extent to which the robustness of the Dl system may be inherent to the Gaussian shape of the Dl gradient, versus how much of the robustness requires a special mechanism, we analyzed an empirical, dosage-scaling description of the gradient.

Let *c*(*x*) be the distribution of nuclear Dl as a function of the DV coordinate *x* :

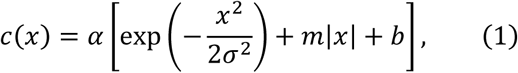

where *α* is a proportionality constant related to morphogen dosage, *σ* represents the spatial width of the Dl gradient, *m* is the shallow, downward slope of the Dl gradient tail, and *b* represents the basal levels of the gradient, related to the levels of Dl that is present in the dorsal-most nuclei. From empirical measurements, *b* ≈ 0.4 and *m* ≈ −0.1 [20].

To calculate the robustness of the predicted gene expression boundaries with respect to changes in *α*, we performed a sensitivity analysis. Let the sensitivity coefficient of a gene expression border with respect to maternal *dl* dosage be defined as *ϕ* ≡ (*∂* ln *x_g_*/*∂* ln *α*), where *x_g_* is the location of gene expression boundary and *θ* is the threshold in Dl nuclear concentration required to express the gene (see Materials and Methods for more information). We found the model of the Dl gradient described by Eqn (1) has unacceptably high values of the sensitivity coefficient (**Fig. 2A**). As a rule of thumb, sensitivity coefficients should be designed to be 0.3 or less [32, 33]; the gradient described by Eqn (1) has sensitivity coefficients of one or greater.

**Figure 2:**
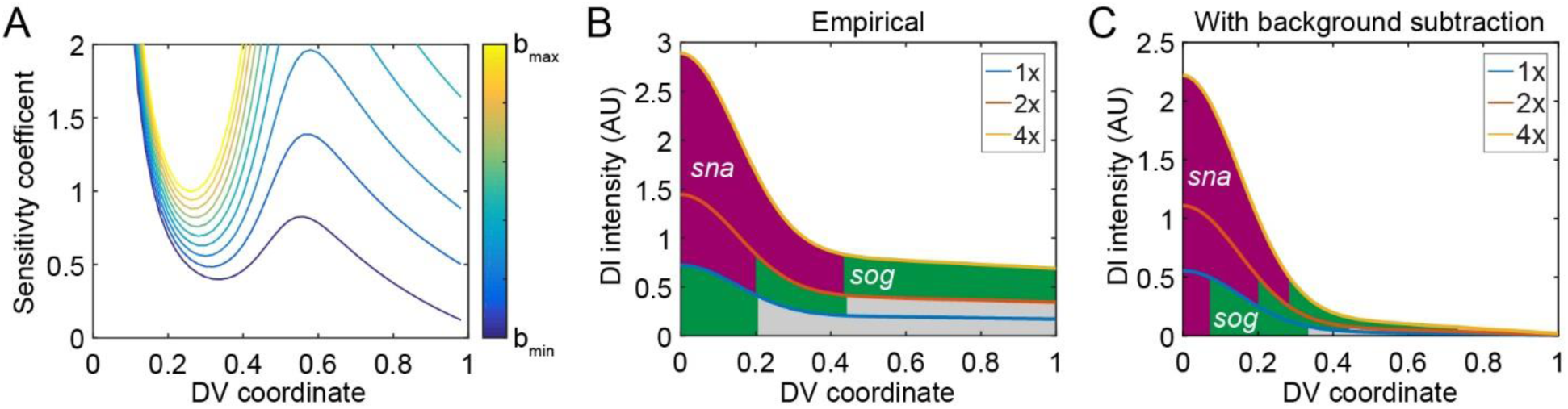
Theoretical consideration of the sensitivity coefficient. (A) Testing whether a lower value of the parameter b could result in a lower sensitivity. (B) The empirical prediction shows that 1x embryos completely lose *sna* expression, while 4x embryos have an overexpanded domain of *sna*, and lose *dpp* completely. (C) The prediction when lower b values were used.

Previously, it was found that a model in which both Dl and Dl/Cact complex are present in the nucleus was more consistent with experimental results than one in which only free Dl is allowed to enter the nucleus [22]. This model was also more robust to noise in Dl levels. Therefore, we asked whether empirically modeling the presence of Dl/Cact complex in the nuclei could also improve the predicted robustness with respect to maternal *dl* dosage. In this case, Eqn (1) represents the sum of the two Dl-containing species. Deconvolution of the non-functional Dl/Cact contribution from the sum would result in the active, “true” Dl gradient (i.e., free Dl). Our previous work has suggested the Dl/Cact contribution is roughly constant across the DV axis [22], so that empirically, the active Dl gradient can be modeled by Eqn (1) with a much lower value of *b*. If we set *b* = 0.11, so that the intensity of free nuclear Dl in the dorsal-most nuclei is 1% of the intensity in the ventral-most nuclei, the sensitivity of gene expression is improved markedly (Fig. 2A). However, even in this best-case scenario, the minimum sensitivity coefficient (located at *x_g_* = 0.34) remains unacceptably high, roughly 0.4, while gene expression boundaries located elsewhere experience even higher sensitivities.

To put the problem in more experimentally concrete terms, we can use Eqn (1) to predict the outcome of deleting one copy of maternal *dl* (1x *dl*), or expressing two extra copies (4x *dl*). Let *α* = 1 to represent the wildtype dosage of maternal *dl*, so that *α* = 0.5 and *α* = 2 represent the 1x and 4x embryos, respectively. In the perturbed cases, the predicted DV gene expression profile in the embryo would result in lethality: 1x embryos completely lose *sna* expression, while 4x embryos have a highly expanded domain of *sna* and lose *dpp* completely (Fig. 2B). As with the sensitivity coefficient above, if *b* is lowered, the effects on gene expression are less severe (Fig. 2C). However, the empirical model still predicts lethality: 1x embryos express *sna* in < 10% of the DV axis [26], and 4x embryos have severely reduced *dpp* expression. We conclude that robustness does not arise simply from a Gaussian shape in a dosage-scaling context, and thus, there must be a mechanism by which the embryo compensates for changes in the maternal *dl* dosage.

### Robustness of Dl-dependent gene expression

While the dosage-scaling model predicted high sensitivity of gene expression, limited measurements of gene expression in 1x and 4x embryos [5, 26], as well as their viability [27–29], suggest the system is robust. To more accurately quantify the robustness of Dl target gene expression with respect to *dl* dosage, we performed large sample size measurements (generally n ∼ 40 or greater) of the expression of two Dl target genes, *sna* and *sog*, in 1x, 2x, and 4x embryos. We found that, with only one exception, the expression domains of both genes in 1x and 4x embryos were statistically different from their expression in 2x (wildtype) embryos (p-val ≤ 2× 10^−4^; Fig 3). The lone exception, the *sna* border in 4x embryos, had a much smaller sample size than the rest (n = 13). Furthermore, the direction of the shifts in gene expression boundaries were as one might expect: in 1x embryos, the gene expression domains shifted closer to the ventral midline, while in 4x embryos, they shifted more dorsally.

**Figure 3:**
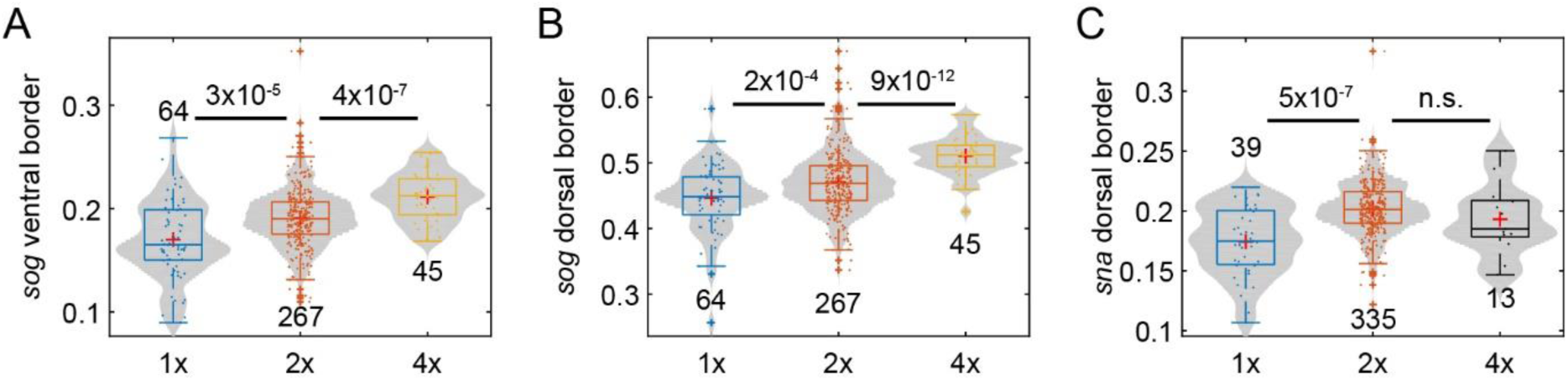
Varying the maternal *dl* dose influences gene expression. (A) Box-and-violin plot of the ventral border of *sog*. (B) Box-and-violin plot of the dorsal border of *sog*. (C) Box-and-violin plot of the of the dorsal border of *sna*. The numbers above or below distributions indicate sample size Numbers between distributions indicate p-value; n.s. = “not significant”. Plus signs indicate statistical outliers.

Even though we were able to measure statistically significant differences from wildtype, the shifts in gene expression borders were minimal (roughly 10% or less; see Table 1), in contrast to the predictions of the dosage-scaling model (Eqn 1). Therefore, our quantification of the robustness of gene expression domains further suggests that a mechanism exists to mitigate the effects of altering the dosage of maternal *dl*. We suspected that this mechanism could be traced to the shape and width of the Dl gradient in 1x, 2x, and 4x embryos.

**Table 1:** Average gene expression locations or Dl gradient widths in 1x, 2x, and 4x embryos. The percent columns are the absolute percent change from wildtype.

| Property | 1x | 1x (%) | 2x (wt) | 4x | 4x (%) |
| --- | --- | --- | --- | --- | --- |
| *sna* boundary | 0.17 | 14 | 0.2 | 0.19 | 4 |
| *sog* ventral boundary | 0.17 | 11 | 0.19 | 0.21 | 10 |
| *sog* dorsal boundary | 0.45 | 5 | 0.47 | 0.51 | 8 |
| DI gradient width | 0.13 | 15 | 0.15 | 0.17 | 11 |

### Robustness of the Dl gradient

Previously, it has been shown that a half maternal dose of *dl* significantly shortens and widens the Dl gradient, and results in a flattened, and sometimes double-peaked, top [5,24–26]. Given that this outcome cannot be predicted by the non-robust dosage-scaling description, in which the width and shape of the gradient do not change with dosage, we asked whether such changes to the Dl gradient would be sufficient to confer robustness to predicted gene expression. Therefore, we measured the Dl gradient in embryos loaded with 1x, 2x, and 4x copies of maternal *dl* (see Fig. 4A and Table 1). As previously reported, 1x embryos had a wider and flatter Dl gradient [5, 26]. However, the previously-reported width measurements for 1x embryos cannot be directly compared to widths in wildtype embryos, given the width measurements are based on the assumption that the Dl gradient is Gaussian-shaped, which the 1x Dl gradient is not. Accounting for the differing shape (see Supplementary information), the 1x Dl gradient measures as narrower than wildtype (Fig. 4B).

**Figure 4:**
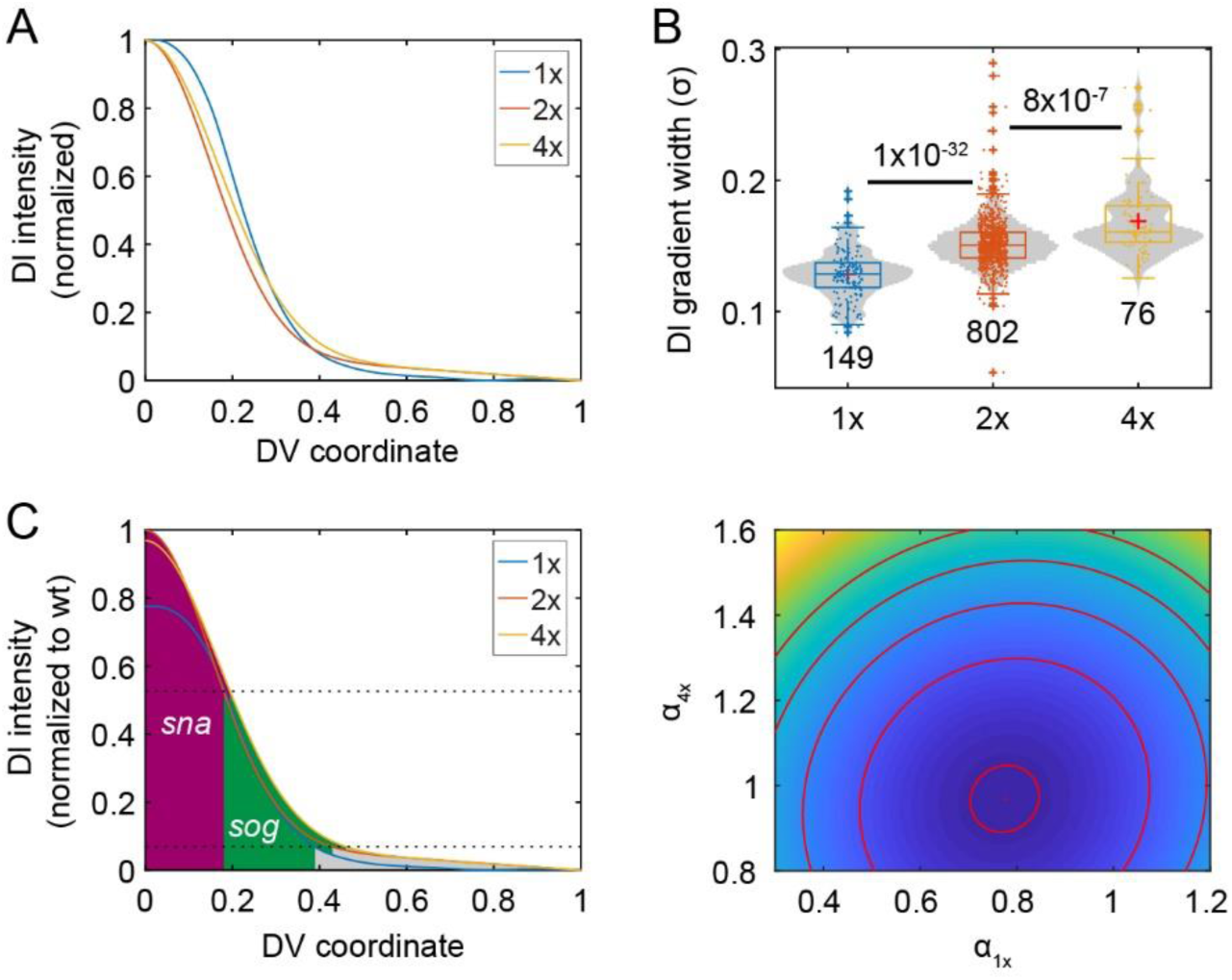
Varying the maternal *dl* dose influences the Dl gradient. (A) Averaged and normalized Dl gradients in 1x, 2x, and 4x embryos. Averaged from n > 10 embryos. (B) Box-and-violin plot of the width of the Dorsal gradient in the genotypes shown in (A). Numbers below distributions indicate sample size. Numbers above indicate p-values. (C) Graph of Dl gradients with best-fit amplitudes for the 1x and 4x gradients, with respect to the 2x gradient set to amplitude of one. (D) Contour plot of the SSE with respect to the amplitude of the 1x gradient (*α*_1*x*_) and that of the 4x gradient *α*_4*x*_. Red dot: the set of best-fit amplitudes. Red curves show the contours of the objective function landscape.

When we examined 4x embryos, we found the gradient became statistically wider (Table 1; Figs. 4 A,B), which also defies a dosage-scaling description of the Dl gradient. However, rather than explaining the robustness of the Dl system, these measurements naïvely predict even higher sensitivities than the dosage-scaling model. Consider the basic expectation that the 1x gradient should have a roughly 50% lower amplitude than wildtype, while the 4x gradient should have a roughly 200% higher amplitude, even if the gradients are not the exact shape and width as wildtype. The combination of decrease in gradient amplitude and decrease in gradient width in 1x embryos, or an increase of both in 4x embryos, would likely result in sensitive Dl-dependent gene expression, as the two effects (amplitude and width) exacerbate each other. In contrast, the dosage-scaling model has only one effect: a changing gradient amplitude.

One way to explain the robustness of gene expression, given the observed changes in Dl gradient shape and width, would be if the amplitudes of the 1x and 4x gradients significantly departed from expectation. Therefore, we computed the amplitudes for the 1x and 4x gradients, with respect to the 2x gradient (which was set to an amplitude of one), that would most closely predict the experimentally observed gene expression in these embryos (Fig. 4C; see Supplementary Methods). We found that a 1x gradient with > 50% of the wildtype amplitude, and a 4x gradient with < 100% of the wildtype amplitude, would be consistent with the experimentally observed robust gene expression. While it is unlikely the 4x gradient would have a shorter amplitude than the 2x gradient, values slightly greater than one are also acceptable (Fig. 4D).

These results suggest that the mechanism to impart robustness with respect to morphogen dosage can control the width, shape, and amplitude of the Dl gradient. Our previous work has shown that facilitated diffusion, also known as shuttling, combined with saturation of the active Toll receptor, can produce the wider, flatter gradients observed in the 1x embryos [26]. Given the saturation of the active Toll receptor, this same mechanism may allow for negligibly taller Dl gradients in 4x embryos. Furthermore, we have seen that 1x embryos that also have compromised shuttling are non-viable [26]. Therefore, to test whether such a combination of mechanisms – shuttling and Toll saturation, together with deconvolution (Fig. 2C) – can grant the Dl gradient system robustness with respect to maternal *dl* dosage, we analyzed a mechanistic model capable of capturing these mechanisms.

### Computational modeling of Dl gradient sensitivity

The model of Dl/Cact interactions analyzed here is based on previous models of the Dl gradient [22,24,26,30]. In particular, we assume that Dl, Cact, and Dl/Cact complex can bind, diffuse, and enter and exit the nucleus; and that Toll signaling can be modeled using a Michaelis-Menten-like formulation (see Methods and Eqns 2-4) [26]. Using this model, we performed a random parameter search to screen for parameter sets in which the Dl nuclear gradient was robust to changes in maternal *dl* dosage (Fig. 5 A-G, see Methods for more details).

**Figure 5:**
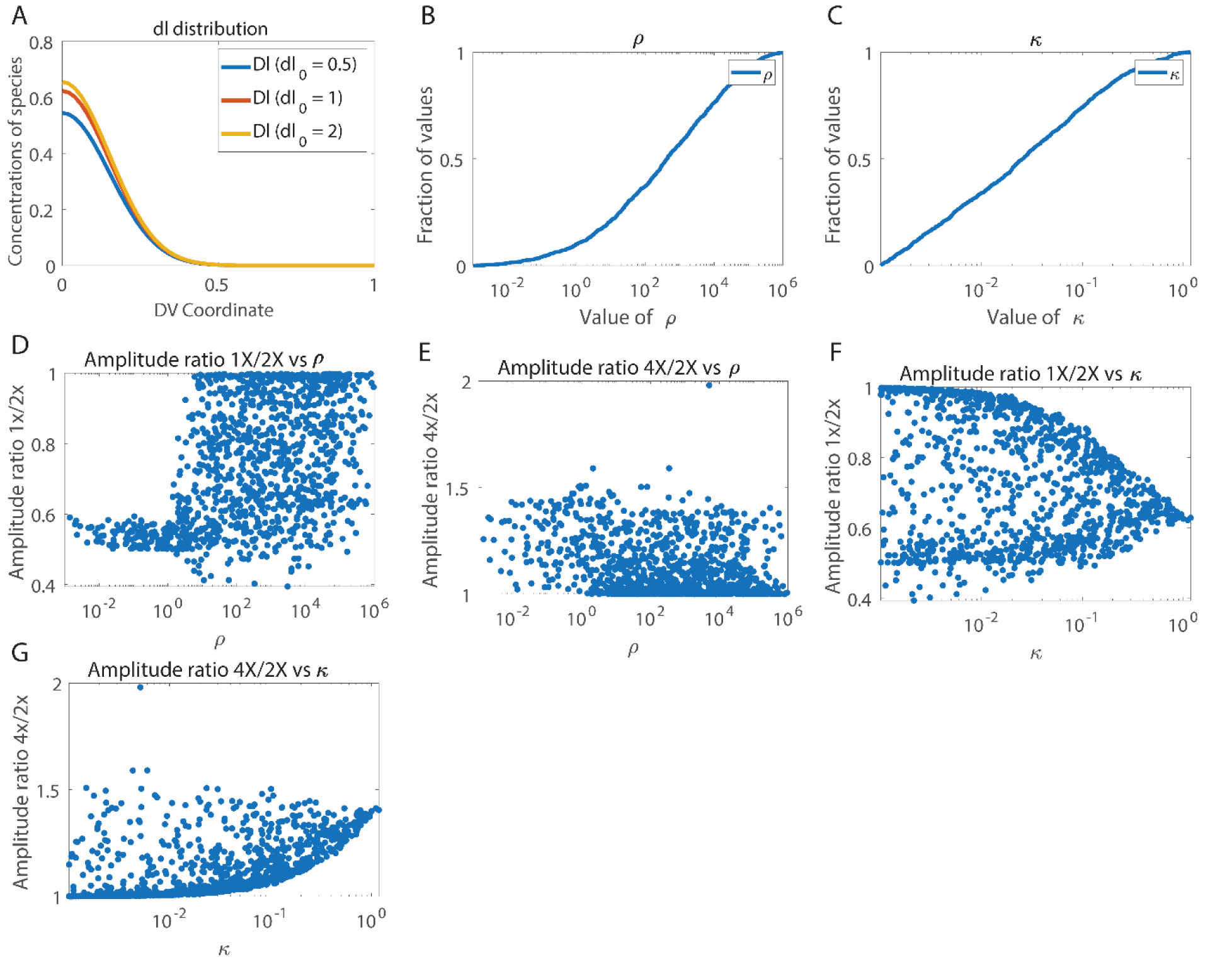
Computational results. (A). Concentration distribution of free Dl for one of the robust parameter sets for dosage 1x,2x and 4x. (B). Cumulative distribution plot for length scale ratio (ρ) (C). Cumulative distribution plot of the Michaelis Menten constant (κ) (D). Plot of Amplitude ratio 1x/2x against length scale ratio (E). Plot of Amplitude ration 2x/4x against length scale ratio (F). Plot of Amplitude ratio 1x/2x against κ (G). Plot of length scale ratio 2x/4x against κ.

We found three trends in the parameter sets necessary for robustness. First, all robust parameter sets predicted that the free Dl nuclear intensity drops to near zero on the dorsal side of the embryo, a result consistent with the deconvolution hypothesis that suggests that Dl fluorescence, as observed in immunostaining experiments or in live embryos expressing Dl-GFP, represents both free Dl and Dl/Cact complex, and that it is important to distinguish between the two [22]. This observation may be similar to the result seen in Fig. 2C, in which decay of the Dl gradient to zero at the dorsal midline improved robustness. Fig. 5A shows the concentration gradient of free Dl for all three values of dosage (1x, 2x, 4x) for one of the robust parameter sets; it can be seen that all concentration curves fall to zero around lateral regions of the embryo. These same parameter sets predict a gradient of Dl/Cact complex that is non-zero at the dorsal midline (S1 Fig), suggesting that, in these simulated embryos, direct fluorescence measurements (the sum of Dl and Dl/Cact in the nucleus) would reveal what appears to be a non-robust Dl gradient. Furthermore, the model does not universally predict that the Dl gradient decays to zero at the dorsal midline. While all robust parameter sets do so, many rejected parameter sets do not (S2 Fig). Thus, the model results strongly suggest deconvolution is necessary for robustness.

Second, we found that the effective diffusivity of Dl/Cact complex is greater than that of free Dl in nearly all robust parameter sets. As the flux of Dl/Cact complex is ventrally directed (i.e., shuttling), this result implies that there is a net flux of Dl from dorsal to ventral regions [26]. We plotted the distribution of the ratio of the effective diffusivity of Dl/Cact complex to that of free Dl, henceforth called ratio of length scales (ρ), in Fig. 5B. In over 95% of the robust parameter sets, this ratio was found to be greater than one. Thus, the constraint of robust gene expression rejects most parameter sets that do not entail facilitated diffusion of Dl by Cact.

Finally, the model also suggests that saturation of Toll receptors is necessary for robustness of gene expression. In all robust parameter sets, the saturation constant for Toll signaling, *κ*, was between 0.001 and 2. Indeed, as seen in Fig.5C, *κ* was the most tightly constrained parameter, which implies tight regulation of Toll saturation may be the most important aspect of the mechanism to ensure robustness of gene expression. This shows that the constraints in the model overwhelmingly favor saturation of Toll receptors by the Dl/Cact complex, as the concentration of Dl/Cact complex in our equations has been scaled to be of order 1. Taken together, these modeling results suggest that the mechanisms of deconvolution and Toll saturation are necessary for a robust DV system, while shuttling of Dl by Cact greatly improves the chances of robustness [22, 26].

### Model predictions of amplitude ratios

In addition to endorsement of the above three mechanisms, the model makes a specific prediction regarding the ratios of amplitudes of the 1x and 4x embryos to that of the wildtype (2x). As seen in Fig. 5 D,E the ratio of amplitudes of 1x embryos:2x embryos is overwhelmingly favored to be greater than 0.5 but less than 0.9, and that of 4x embryos:2x embryos is favored to fall between 1 and 1.55.

It can be seen from Fig. 5 D,E that as the value of *κ* decreases (and thus, Toll becomes more saturated), the range of amplitude ratios available to robust descriptions of the Dl gradient increases. For lower values of *κ*, a range of 1.1 to 1.6 is accepted by the model for ratio of amplitudes of 4x embryos:2x embryos and a range of 0.5 to 0.85 for ratio of amplitudes of 1x embryos:2x embryos. At higher values of *κ*, the values of the amplitude ratios for 4x/2x and 1x/2x seem to converge to 1.45 and 0.57 respectively, meaning that as Toll is more easily saturated (lower values of *κ*), the model allows for a small, but noticeable range of amplitude ratios for varying dosages. This result seems to indicate that under constrained conditions of Toll saturation, only particular peak amplitudes are preferred – about 1.55 times the wildtype value for 4x embryos and about 0.57 times the wildtype value for 1x embryos. However, if Toll receptors saturate easily, an appreciable range of amplitude ratios leads to robustness. Thus, Toll saturation seems to be an inherent mechanism for robustness in the embryo.

In a similar way, the model predicts that the extent of shuttling of Dl by Cact also affects the acceptable values of the amplitude ratios. In Fig. 5 F,G we plotted amplitude ratios against ratio of length scales, and we see that when facilitated diffusion by Cact does not occur (about 5% of parameter sets), the amplitude ratios of 4x/2x and 1x/2x are constrained around 1.45 and 0.57, respectively. The values of amplitudes seem to converge to similar values when Toll saturation was minimal. On the other hand, when the length scale ratio is greater than 1, a wider range of amplitude ratios are accessible to the embryo.

Thus, it seems that both Toll saturation and shuttling of Dl from dorsal to ventral regions allows the embryos to explore a wider range of amplitude ratios, which allows greater flexibility for robustness. However, when the above mechanisms are constrained, the amplitude ratios must take on specific values, which in turn makes it difficult to achieve robustness.

## Discussion

Animal development is a complex process that must be buffered against myriad environmental, nutritional, and genetic perturbations. The robustness of development with respect to these perturbations often requires regulatory mechanisms. Here we investigated the robustness of gene expression in the early *Drosophila* embryo with respect to variations in the maternal gene dosage of the NF-κB transcription factor Dorsal in a quantitative and computational manner. The NF-κB pathway is highly conserved and is centrally involved in a diverse array of cellular processes, including inflammation, apoptosis, and innate immunity. In flies, Dl/NF-κB also directs embryonic development and differentiation. However, essential questions related to NF-κB robustness in *Drosophila* remain unresolved. Our analysis of an empirical, dosage-scaling description of the Dl gradient, together with detailed measurements of the Dl gradient and its target genes, suggest that a mechanism to control the shape, width, and amplitude of the Dl gradient is necessary for robustness. Our previous work found three novel mechanisms in the establishment of the Dl gradient: deconvolution, shuttling, and Toll saturation [22, 26]. In this paper, we used a computational model to study the importance of each of these mechanisms for the robustness of the Dl system.

Recent work showed the importance of deconvolving experimentally-measured fluorescence signal into free Dl and bound Dl (Dl/Cact complex) when interpreting the Dl gradient [22]. Doing so results in a nuclear Dl gradient that drops to near zero instead of to non-zero basal levels at dorsal regions [5,15,20,21]. In the dosage-scaling model, deconvolution was modeled by setting basal levels to near zero. While this choice of basal levels improved robustness somewhat in the dosage-scaling model, the gene expression boundaries remained overly sensitive to *dl* dosage. Furthermore, every robust parameter set in the computational model predicted a free Dl gradient that decayed to near zero, whereas non-robust parameter sets did not. Thus, while deconvolution by itself is not sufficient to explain the robustness of gene expression boundaries, it appears to be a necessary piece.

Our model also suggests the shuttling mechanism increases robustness of the Dl system. In such a mechanism, Toll signaling creates a sink for Dl/Cact complex, which establishes a ventrally-directed flux to accumulate Dl in ventral regions. While it is possible that free Dl then diffuses dorsally, such counter-diffusion is likely mitigated by capture of free Dl by the nuclei. Previous work in our lab suggests that shuttling of Dl/Cact complex from dorsal to ventral regions is an important factor for robustness in the embryo [26]. Our model supports this result, as most parameter sets that selected for robust gene expression favored facilitated diffusion of Dl by Cact, as the effective diffusivity of Dl/Cact was higher than that of free Dl.

Previous work also suggested that, in wildtype embryos, active Toll receptors are limiting [26], thereby maintaining robust gene expression, even when *dl* dosage varies from wildtype. In wildtype embryos, when active Toll signaling complexes are saturated with Dl/Cact complex, a significant number of Dl/Cact complexes bypass the ventral-lateral regions without being dissociated, and Dl is shuttled to the ventral-most portions of the embryo. On the other hand, if active Toll signaling complexes are not saturated, as may be the case in 1x embryos, the Dl/Cact complex will be dissociated at a higher rate in the ventral-lateral regions of the embryo and will be unable to reach the ventral-most regions of the embryo. The lack of Toll saturation in 1x embryos thus results in a flatter and wider concentration gradient of nuclear Dl.

While embryos with 4 copies of dl have double the wildtype Dl dose, twice as much Dl will not necessarily enter the nuclei because that process relies on Toll signaling, which may be saturated. Similarly, decreasing the *dl* dosage, as in the case of 1x embryos, implies halving the amount of Dl/Cact complex without reducing the absolute number of free Dl molecules that will enter the nuclei. If the active Toll complexes remain constant in all three cases of dosage and provided that they are saturated, only minor variations occur in the concentration of free Dl when dosage changes (Fig 5A).

In this work we have demonstrated the importance of certain built-in mechanisms within the early *Drosophila* embryo that ensure robustness of gene expression along the DV axis. These three mechanisms, (deconvolution of the measured Dl fluorescence into free Dl and Dl/Cact complex, saturation of Toll receptors by Dl/Cact complex, and shuttling of Dl by Cact from dorsal to ventral regions of the embryo) are crucial for ensuring that genes expressed in the DV axis have domain boundaries in specific regions. We have presented both experimental and computational evidence that these processes are paramount for safeguarding against genetic perturbations to *dl* dosage. The advances in studying the molecular mechanism behind robustness with respect to maternal *dl* dosage may open the door for understanding the question of how sustained embryonic development can be achieved despite genetic and environmental fluctuations.

## Methods

### Fluorescent in situ Hybridization

All embryos were aged to NC 14 (approx. 2-4 hours after egg lay), then fixed in 37% formaldehyde according to standard protocols [34]. A combination fluorescent *in situ* hybridization/fluorescent immnuostaining was performed according to standard protocols [34]. Briefly, fixed embryos were washed in PBS/Tween and hybridized at 55 °C overnight with anti-sense RNA probes, which were generated according to standard lab protocol. The embryos were then washed and incubated with primary antibodies at 4 °C overnight. The next day, they were washed and incubated for 1-2 hrs with fluorescent secondary antibodies at room temperature. The embryos were then washed and stored in 70% glycerol at −20 °C. Embryos were imaged within one month of completing the protocol.

Antibodies used were anti-dorsal 7A4 (deposited to the DSHB by Ruth Steward (DSHB Hybridoma Product anti-dorsal 7A4)) (1:10), donkey anti-mouse-488 (Invitrogen A21202, Lot 81493) (1:500), rabbit anti-histone (abcam ab1791, Lot 940487) (1:5000), donkey anti-rabbit-546 (Invitrogen A10040, Lot 107388) (1:500), goat anti-biotin (ImmunoReagents, Raleigh, NC, GtxOt-070-D, Lot 19-19-112311) (1:50,000), donkey anti-goat-647 (Invitrogen A21447, Lot 774898) ((1:500), goat anti-fluorescin (Rockland 600-101-096, Lot 19458) (1:500), rabbit anti-fluorescin (Life Technologies A889, Lot 1458646) (1:500), goat anti-histone (Abcam, ab12079, Lots GR6952-4 and GR129411-1) (1:100), donkey anti-rabbit-350 (ImmunoReagents, DkxRb-003-D350NHSX) (1:500). For some experiments the nuclear stain Draq5 (Cell Signaling #4084S) was used instead of an anti-histone antibody.

### Mounting and Imaging of Fixed Embryos

Embryos were cross sectioned and mounted in 70% glycerol as described previously [35]. Briefly, a razor blade was used to remove the anterior and posterior thirds of the embryo, leaving a cross section roughly 200 µm long by 200 µm in diameter. These sections were then oriented such that the cut sides became the top and bottom. Sections were then imaged at 20x on a Zeiss LSM 710 microscope. 15 z-slices 1.5 µm apart were analyzed, for a total section size of 21 µm.

### Image analysis

Images of embryo cross sections were analyzed using a previously derived algorithm [36]. Briefly, the border of the embryo was found computationally, then the nuclei were segmented using a local thresholding protocol. The intensity of dl in each segmented nucleus was calculated as the ratio between the intensity in the dl channel divided by the intensity in the nuclear channel. The intensity of mRNA expression was calculated as average intensity within an annulus roughly 18 μm wide around the perimeter of the embryo. mRNA profiles were fit to canonical parameters; those with a goodness of fit (gof) less than 0.7 were omitted from study.

All dl gradients were fit to a Gaussian, and these fits were used to determine the width parameter, σ. Gradients with a gof less than 0.8 were eliminated from the results. Normalized intensity plots were generated by fitting each embryo’s data to its own Gaussian by subtracting the B value and 70% of the M value, then dividing by the A value. (X = (x – B – 0.7M)/A)). The average normalized intensity plot was generated by averaging the plots of all embryos in the specified genotype.

Multiple experiments with statistically similar wild type controls were analyzed simultaneously to generate the data in this report. Statistical significance was calculated using two-tailed homoscedastic t-tests.

### Model equations

The equations for the computational model are as follows:

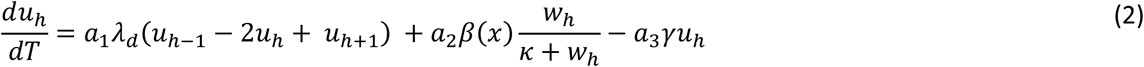

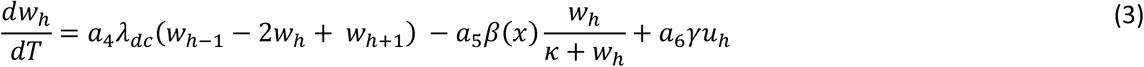

Where 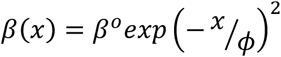 represents the gaussian Toll-mediated rate constant and *κ* represents the Michaelis Menten constant for the dissociation of Dl/Cact complex; *u* and *w* represent cytoplasmic species Dl and Dl/Cact complex respectively; subscript *h* represents a nucleus and its associated cytoplasmic compartment; *λ_i_* represents effective intercompartmental exchange rates; and the *α_i_*’s are weighting factors related to the nuclear import/export equilibrium constants and the geometry of the nucleus and cytoplasm (see Supplementary Information for more details).

Equations (2-3) above have been derived after simplifying a more detailed model (see Supplementary information for details). The nuclei are modeled as spheres sitting in cuboidal cytoplasmic compartments that span the periphery of the embryo. Since the embryo is approximately symmetric about the DV axis; the spatial coordinate was varied from 0 to 1 with the former representing the ventral midline and the latter, the dorsal midline. The number of such compartments/nuclei/cells is taken to be 51, approximately equal to the number of nuclei in NC 14 found from live fluorescence imaging [20]. Both nuclei and the cytoplasm volumes are considered well mixed. We assume that the nucleus and cytoplasm are in a state of pseudo-equilibrium. Thus, *k_out_C_nuc_* ≈ *k_in_C_cyt_* or *C_nuc_* ≈ *K_eq_C_cyt_* where, *K_eq_* ≡ *k_in_*/*k_out_* is defined as the equilibrium constant for nuclear import/export for all species. The effect of Toll was modeled with a Michaelis Menten formulation, assuming the concentration of the intermediate species Dl-Cact-Toll to be approximately constant in nuclear cycle 14. The above equations were then non-dimensionalized, approximately with respect to the conditions found in wildtype *Drosophila* embryos at the beginning of NC 14, such that every term was of order 1. The ratio of effective diffusivities or the length scale ratio was then defined as

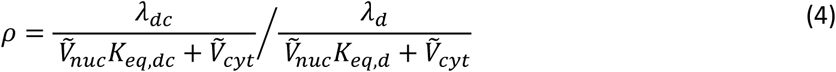

where 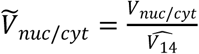 (see Supplementary information for details).

The simulation was run for 60 min, which approximates the time period of NC 14, which is the longest nuclear cycle of the blastoderm. Dosage was varied by doubling or halving the initial concentration of Dl/Cact. The ldimensionless constants obtained from it were then varied from 1e-3 to 1e+3 to obtain concentration profiles for Dl and Dl/Cact. From these concentration profiles, the dorsal border of *sna* and the ventral and dorsal borders of *sog* were calculated assuming the borders are defined by thresholds of free Dl concentration. These model predictions of the borders were compared with experimental values in the least square error sense and parameter sets with errors lower than a set value were accepted as robust (see Supplementary information for details).

## Supporting information

S1 File. This file contains details of methodology.

