## Supplemental Methods and Figures for "Robustness of the Dorsal morphogen gradient with respect to morphogen dosage"

### *Detailed computational model*

Previous models of the Dl nuclear gradient can be classified in terms of their complexity and the number of realistic features they support. In 2009, Kanodia et al. published pioneering modeling work on the Dl gradient, which captured the establishment of Dl gradient by an interaction between Dl, Cact and Toll in the cytoplasm. It was found that this model was inconsistent with the data from live measurements of Venus-tagged Dl. The only way to work around this inconsistency was to assume the presence of nuclear Cact, thus nuclear Dl/Cact. The following model equations represents the full model for the Dl system, which consists of three species namely, Dl, Cact and Dl/Cact complex that are allowed to move between cytoplasmic compartments and between nucleus and cytoplasm within a cell.

$$\frac{d\left[V_{nuc}C_{d,nuc}^{h}\right]}{dt} = A_{nuc}\left(k_{in,d}C_{d,cyt}^{h} - k_{out,d}C_{d,nuc}^{h}\right) - V_{nuc}\left(k_b C_{d,nuc}^{h} C_{c,nuc}^{h}\right) \quad (1)$$

$$\frac{d\left[V_{cyt}C_{d,cyt}^{h}\right]}{dt} = A_{cyt}\Gamma_d\left(C_{d,cyt}^{h-1} - 2C_{d,cyt}^{h} + C_{d,cyt}^{h+1}\right) + V_{cyt}\left(\frac{k_d(x)C_{dc,cyt}^{h}}{\kappa + C_{dc,cyt}^{h}} - k_b C_{d,cyt}^{h} C_{c,cyt}^{h}\right) - A_{nuc}\left(k_{in,d}C_{d,cyt}^{h} - k_{out,d}C_{d,nuc}^{h}\right) \quad (2)$$

$$\frac{d\left[V_{nuc}C_{dc,nuc}^{h}\right]}{dt} = A_{nuc}\left(k_{in,dc}C_{dc,cyt}^{h} - k_{out,dc}C_{dc,nuc}^{h}\right) + V_{nuc}\left(k_b C_{d,nuc}^{h} C_{c,nuc}^{h}\right) \quad (3)$$

$$\frac{d\left[V_{cyt}C_{dc,cyt}^{h}\right]}{dt} = A_{cyt}\Gamma_{dc}\left(C_{dc,cyt}^{h-1} - 2C_{dc,cyt}^{h} + C_{dc,cyt}^{h+1}\right) - V_{cyt}\left(\frac{k_d(x)C_{dc,cyt}^{h}}{\kappa + C_{dc,cyt}^{h}} - k_b C_{d,cyt}^{h} C_{c,cyt}^{h}\right) - A_{nuc}\left(k_{in,dc}C_{dc,cyt}^{h} - k_{out,dc}C_{dc,nuc}^{h}\right) \quad (4)$$

$$\frac{d\left[V_{nuc}C_{c,nuc}^{h}\right]}{dt} = A_{nuc}\left(k_{in,c}C_{c,cyt}^{h} - k_{out,c}C_{c,nuc}^{h}\right) - V_{nuc}\left(k_b C_{d,nuc}^{h} C_{c,nuc}^{h}\right) \quad (5)$$

$$\frac{d\left[V_{cyt}C_{c,cyt}^{h}\right]}{dt} = A_{cyt}\Gamma_c\left(C_{c,cyt}^{h-1} - 2C_{c,cyt}^{h} + C_{c,cyt}^{h+1}\right) + V_{cyt}\left(\frac{k_d(x)C_{dc,cyt}^{h}}{\kappa + C_{dc,cyt}^{h}} - k_b C_{d,cyt}^{h} C_{c,cyt}^{h} - k_{deg}C_{c,cyt}^{h}\right) - A_{nuc}\left(k_{in,c}C_{c,cyt}^{h} - k_{out,c}C_{c,nuc}^{h}\right) + P_c \quad (6)$$

Here, subscripts *nuc* and *cyt* represent nucleus and cytoplasm respectively; *d, c,* and *dc* represent species Dl, Cact, and Dl/Cact complex respectively; superscript *h* represents a nucleus and its associated cytoplasmic compartment. The parameters, $k_{in,species}$ and $k_{out,species}$ represents nuclear import and export rates, $k_b$ represents Dl/Cact binding constant, $\Gamma_{species}$ represents intercompartmental exchange rates, $k_d(x) = k_d^{max}exp\left({}^{x}/_{\phi}\right)^2$ represents the gaussian Toll-mediated rate constant and $\kappa$ represents the Michaelis Menten constant for the dissociation of Dl/Cact complex, $k_{deg}$ represents the degradation rate constant for Cact and $P_c$ represents rate of production of Cact.

The Dl system is represented by 6 equations consisting of Dl, Cact and Dl/Cact in the nucleus and in the cytoplasm. This model is based on previous models used in the literature with some modifications. Firstly, Cact and the Dl/Cact complex were allowed to enter the nucleus and secondly Michaelis Menten kinetics was used to describe the dissociation of the Dl/Cact complex by Toll in the cytoplasm. The width of Toll gradient was fixed at $\phi = 0.15$, which approximates the width of wildtype Dl gradients. In order to minimally describe the effect of dosage of the Dl morphogen on the embryo's development these equations were simplified based on the following assumptions. Firstly, since the time scales of transport of species between adjacent cytoplasmic compartments is much higher than that of nuclear exchange, a state of pseudo equilibrium is assumed between the nucleus and cytoplasm. Thus, $k_{out}C_{nuc} \approx k_{in}C_{cyt}$ or $C_{nuc} \approx K_{eq}C_{cyt}$ where, $K_{eq} \equiv k_{in}/k_{out}$ is defined as the equilibrium constant for nuclear import/export. The values for the equilibrium constants are fixed at $K_{eq,d} = 4$, $K_{eq,dc} = 1$ and $K_{eq,c} = 1$ (1). Secondly, since Cact has a high turnover rate, a uniform concentration of Cact, equal to that at the beginning of nuclear cycle 14 in wildtype embryos, was assumed. Shown below are equations where concentrations have been non-dimensionalized using conditions at the beginning of nuclear cycle 14 in wildtype embryos.

$$\frac{d\left[\left(V_{nuc}K_{eq,d} + V_{cyt}\right)u^h\right]}{dT} = \Gamma_d A_{cyt}\left(u^{h-1} - 2u^h + u^{h+1}\right) + V_{cyt}\left(\frac{\beta(x)w^h}{\kappa + C_d^{wt}w^h} - k_b C_d^{wt} C_c^{wt}\, u^h v^h\right) - V_{nuc}\left(k_b K_{eq,d} K_{eq,c} C_d^{wt} C_c^{wt}\, u^h v^h\right) \tag{7}$$

$$\frac{d\left[\left(V_{nuc}K_{eq,dc} + V_{cyt}\right)w^h\right]}{dT} = \Gamma_{dc} A_{cyt}\left(w^{h-1} - 2w^h + w^{h+1}\right) - V_{cyt}\left(\frac{\beta(x)w^h}{\kappa + C_d^{wt}w^h} - k_b C_c^{wt} u^h v^h\right) + V_{nuc}\left(k_b K_{eq,d} K_{eq,c} C_c^{wt}\, u^h v^h\right) \tag{8}$$

$$\frac{d\left[\left(V_{nuc}K_{eq,c} + V_{cyt}\right)v^h\right]}{dT} = \Gamma_c A_{cyt}\left(v^{h-1} - 2v^h + v^{h+1}\right) - \frac{V_{cyt}}{C_c^{wt}}\left(\frac{\beta(x)C_d^{wt}w^h}{\kappa + C_d^{wt}w^h} - k_b C_d^{wt} C_c^{wt} u^h v^h - k_{deg} C_c^{wt} v^h\right) + \frac{V_{cyt}}{C_c^{wt}}\left(k_b K_{eq,d} K_{eq,c} C_d^{wt} C_c^{wt}\, u^h v^h\right) + \frac{P_c}{C_c^{wt}} \tag{9}$$

where,

$u^h = \frac{C_{d,cyt}^h}{C_d^{wt}}$ $\quad w^h = \frac{C_{dc,cyt}^{wt}}{C_d^{wt}}$ $\quad v^h = \frac{C_{c,cyt}^h}{C_c^{wt}}$ $\quad C_c^{wt} = \frac{P_c}{k_{deg}}$

Due to the high turnover rate of Cact, equation 9, upon non-dimensionalizing simplifies to $v^h = 1$. Finally, the equations in the main text were derived by non-dimensionalizing equations 7 and 8 using the following dimensionless parameters.

$\tilde{V}_{cyt} = \frac{V_{cyt}}{\widehat{V_{14}}}$ $\quad \tilde{A}_{cyt} = \frac{A_{cyt}}{\widehat{A_{14}}}$ $\quad \gamma = -k_b C_c^o \bar{T}$ $\quad \beta = k_d^{max}\bar{T}$ $\quad \lambda_d = \frac{A_{nuc}^{14}\Gamma_d T}{V_{nuc}^{14}}$ $\quad \lambda_{dc} = \frac{A_{nuc}^{14}\Gamma_{dc} T}{V_{nuc}^{14}}$

Thus, based on the two assumptions, the full six equation model was reduced to the two-equation model as shown in the main text.

*Least squares method for determining robustness in the model*

The error in the predictions of boundaries of gene expression was defined, for every border, as follows,

$$e_{\beta}(\theta) = \left(\varepsilon_{\beta,1x}\right)^2 + \left(\varepsilon_{\beta,2x}\right)^2 + \left(\varepsilon_{\beta,4x}\right)^2$$

$$= \left(\frac{x_{\beta,model,1x}(\theta) - x_{\beta,exp,1x}}{\sigma_{\beta,exp,1x}}\right)^2 + \left(\frac{x_{\beta,model,2x}(\theta) - x_{\beta,exp,2x}}{\sigma_{\beta,exp,2x}}\right)^2 + \left(\frac{x_{\beta,model,4x}(\theta) - x_{\beta,exp,4x}}{\sigma_{\beta,exp,4x}}\right)^2 \qquad (10)$$

where, $x_{\beta,model,g}$ is the model boundary prediction,$x_{\beta,exp,g}$ is the experimental measure of border and $\sigma_{\beta,exp,g}$ is the experimentally observed variation in boundary of gene $\beta$ of genotype g.

For any gene expression border $\beta \in B$, where $B$ = {sna, sogd, sogv} and genotype $g \in G$ where $G$ = {1x, 2x, 4x}, the error is calculated by minimizing $e_{\beta}(\theta)$ with respect to its concentration threshold $\theta$. Those parameter sets with error values less than 1.5 for all gene expression boundaries, were deemed robust.

*Approximate gradient width for dl 1x gradients*

As the Dl gradient in embryos from mothers heterozygous for *dl* is not Gaussian-shaped, fitting it to a Gaussian gives an aberrant value for $\sigma$. To attempt to characterize the flat-topped gradients by a value of $\sigma$ equivalent to its closest approximation to a wildtype gradient, we did the following. First, by averaging ~75 Dl gradients from 1x embryos, we created a “canonical” flat-topped gradient, normalized between zero and one, denoted $f_{50}(x)$. Next, we fit each 1x Dl gradient to this canonical gradient by allowing the spatial coordinate to be stretched (see Carrell et al., 2017; Liberman et al., 2009; Trisnadi et al., 2013 for examples). Therefore, for each 1x embryo $i$, we obtained a best-fit value of the spatial stretching factor, $\delta_i$.

Next, we calculated the area under the curve of a wt Gaussian:

$$I_{100} = \int_0^1 \exp\left(-\frac{x^2}{2\sigma^2}\right) dx \approx \int_0^\infty \exp\left(-\frac{x^2}{2\sigma^2}\right) dx = \sigma\sqrt{2}\int_0^\infty \exp(-z^2)\, dz = \sigma\sqrt{\frac{\pi}{2}} \tag{10}$$

where $z = x/(\sigma\sqrt{2})$, and the change of the upper limit of integration to ∞ is valid because $\sigma \leq 0.3$. The average width of the wildtype gradient is $\sigma_{wt} = 0.152$, which implies $I_{100} = 0.1880$.

Next, we calculated the area under the curve of $f_{50}(x)$, which was $I_{50} = 0.2438$. Next, we computed the value of $\alpha_{50}$ makes $\alpha_{50} I_{50} = 0.5 I_{100}$, and found that $\alpha_{50} = 0.3855$. Finally, to calculate the equivalent Gaussian-like width of the 1x Dl gradients, we computed the value of sigma that minimizes the following:

$$\varepsilon = \int_{x_1}^{x_2} [f_{100}(x;\sigma) - \alpha_{50} f_{50}(x)]^2 dx \tag{11}$$

This value of $\sigma$, which we will call $\sigma_{1\times}^{eff}$ is 0.1283. In other words, if the average 1x embryo has 50% of the Dl in an average wildtype embryo, then the Dl gradient in an average 1x embryo looks most like a wildtype gradient with a width of 0.1283 (slightly narrower than the average wildtype gradient). Taking this base value of $\sigma_{1\times}^{eff}$, we can find the effective gradient width for each embryo $i$ by multiplying by $\delta_i$.

*Least squares calculations for thresholds and amplitudes in the empirical description*

To estimate the necessary amplitude of the 1x and 4x canonical curves, with respect to wt, in order to achieve the observed gene expression (and given the observed shape and width of the Dl gradient), we constructed a least squares estimation. Let the objective function $f$ be the sum of the squares of error between the (empirical) Dl gradient at the locations of a given gene expression boundary and the estimated threshold for that gene:

$$f(\boldsymbol{\alpha},\boldsymbol{\theta},\boldsymbol{X},\boldsymbol{S}) = \sum_{g\in G}\sum_{\beta\in B}\left(\varepsilon_{\beta,g}\right)^2 = \sum_{g\in G}\sum_{\beta\in B}\left(\frac{\alpha_g Dl_g\left(x_{\beta,g},\sigma_g\right)-\theta_\beta}{s_{\beta,g}}\right)^2 \tag{12}$$

…where the vector $\boldsymbol{\alpha} = [\alpha_{1x},\alpha_{2x},\alpha_{4x}]$, the vector $\boldsymbol{\theta} = \left[\theta_{sna},\theta_{sogv},\theta_{sogd}\right]$, the set of genotypes is $G = \{1\text{x},2\text{x},4\text{x}\}$, the set of boundaries is $B = \{\text{sna},\text{sogv},\text{sogd}\}$, and $x_{\beta,g}$ is the boundary location and $s_{\beta,g}$ is a measure of the variability for that genotype and boundary. In addition, the position array $\boldsymbol{X}$ and standard error array $\boldsymbol{S}$ are:

$$\boldsymbol{X} = \begin{bmatrix} x_{sna,1x} & x_{sna,2x} & x_{sna,4x} \\ x_{sogv,1x} & x_{sogv,2x} & x_{sogv,4x} \\ x_{sogd,1x} & x_{sogd,2x} & x_{sogd,4x} \end{bmatrix} \tag{13}$$

$$\boldsymbol{S} = \begin{bmatrix} s_{sna,1x} & s_{sna,2x} & s_{sna,4x} \\ s_{sogv,1x} & s_{sogv,2x} & s_{sogv,4x} \\ s_{sogd,1x} & s_{sogd,2x} & s_{sogd,4x} \end{bmatrix} \tag{14}$$

This can also be written more transparently as:

$$f(\boldsymbol{\alpha},\boldsymbol{\theta},data) = \sum_{\beta\in B}\left(\varepsilon_{\beta,1x}\right)^2 + \left(\varepsilon_{\beta,2x}\right)^2 + \left(\varepsilon_{\beta,4x}\right)^2 \tag{15}$$

$$= \sum_{\beta \in B} \left( \frac{\alpha_{1x} Dl_{1x}\left(x_{\beta,1x}, \sigma_{1x}\right) - \theta_\beta}{s_{\beta,1x}} \right)^2 + \left( \frac{Dl_{wt}\left(x_{\beta,2x}, \sigma_{2x}\right) - \theta_\beta}{s_{\beta,2x}} \right)^2 + \left( \frac{\alpha_{4x} Dl_{wt}\left(x_{\beta,4x}, \sigma_{4x}\right) - \theta_\beta}{s_{\beta,4x}} \right)^2$$

This function can be minimized by least squares, with respect to varying $\alpha_{1x}, \alpha_{4x}, \theta_{sna}, \theta_{sogv}, \theta_{sogd}$.

http://dev.biologists.org/content/develop/144/23/4450.full.pdf

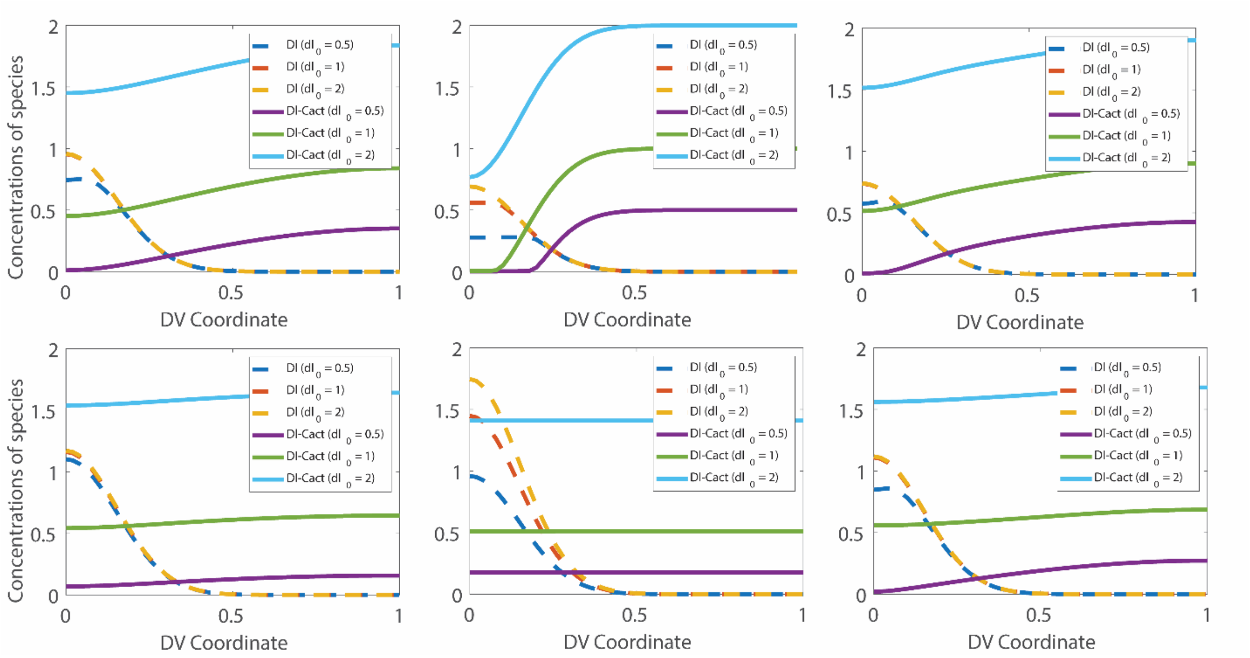


**Figure S1**. Concentration profiles of free Dl and Dl/Cact (robust parameters). This figure shows concentration profiles of free Dl & Dl/Cact, for parameter sets that were accepted as robust. The plots show non-zero concentration for Dl/Cact complexes at the dorsal midline at $x = 1$.

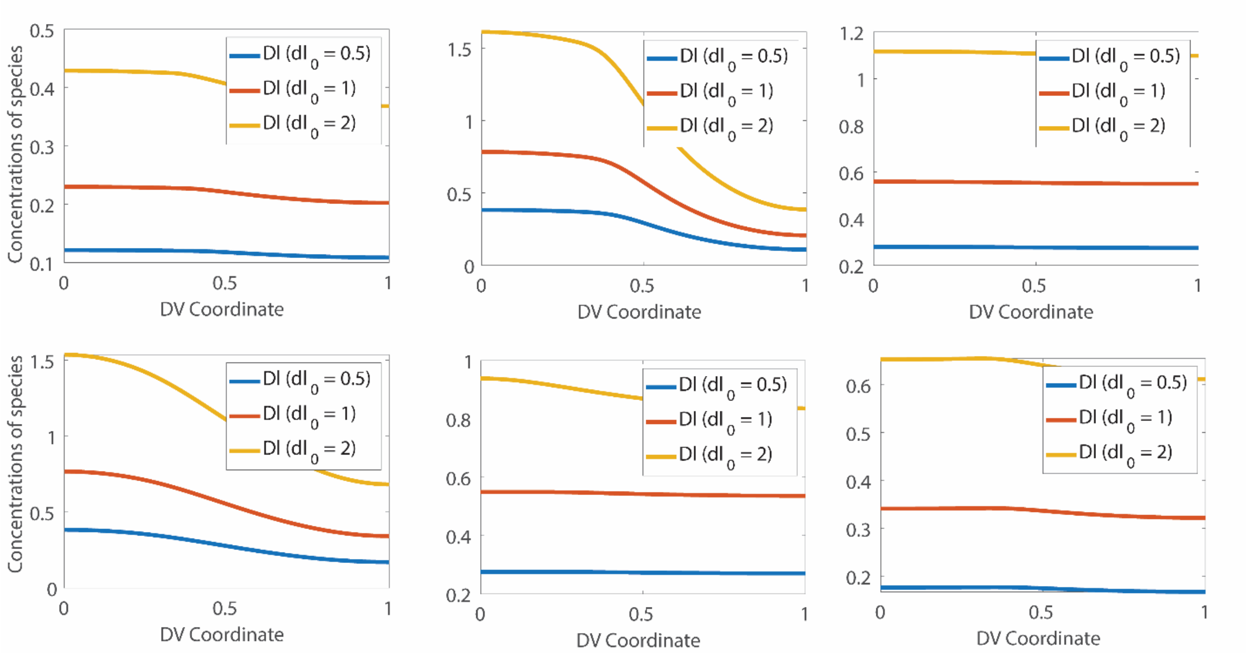


**Figure S2**. Concentration profiles of free Dl (non-robust parameters). This figure shows concentration profiles of free Dl, for parameter sets, that were rejected as not robust. In most cases, concentration curves do not decay to zero at the dorsal midline.
